# Role of prefrontal cortex projections to the nucleus accumbens core in mediating the effects of ceftriaxone on cued cocaine relapse

**DOI:** 10.1101/2020.01.20.911792

**Authors:** Allison R. Bechard, Carly N. Logan, Javier Mesa, Yasmin Padovan Hernandez, Harrison Blount, Virginia L. Hodges, Lori A. Knackstedt

**Author notes:** Corresponding author: Lori A. Knackstedt, University of Florida, 114 Psychology, 945 Center Dr., Gainesville, FL 32611.

## Abstract

Ceftriaxone is an antibiotic that reliably attenuates the reinstatement of cocaine-seeking after extinction while preventing the nucleus accumbens (NA) core glutamate efflux that drives reinstatement. However, when rats undergo abstinence without extinction, ceftriaxone attenuates context-primed relapse but NA core glutamate efflux still increases. Here we sought to determine if the same would occur when relapse is prompted by both context and discrete cues (context+cues) after cocaine abstinence. Male rats self-administered intravenous cocaine for 2 hr/day for 2 weeks. Cocaine delivery was accompanied by drug-associated cues (light+tone). Rats were then placed into abstinence with daily handling but no extinction training for two weeks. Ceftriaxone (200 mg/kg IP) or vehicle was administered during the last 6 days of abstinence. During a context+cue relapse test, microdialysis procedures were conducted. Rats were perfused at the end of the test for later Fos analysis. A separate cohort of rats was infused with the retrograde tracer cholera toxin B in the NA core and underwent the same self-administration and relapse procedures. Ceftriaxone increased baseline glutamate and attenuated both context+cue-primed relapse and NA core glutamate efflux during this test. Ceftriaxone reduced Fos expression in regions sending projections to the NA core (prefrontal cortex, basolateral amygdala, ventral tegmental area) and specifically reduced Fos in prelimbic cortex and not infralimbic cortex neurons projecting to the NA core. Thus, when relapse is primed by drug-associated cues and context, ceftriaxone is able to attenuate relapse by preventing NA core glutamate efflux, likely through reducing activity in prelimbic NA core-projecting neurons.

## Introduction

Cocaine use disorder is associated with relapsing cocaine-seeking even after long periods of abstinence (O’Brien 2003), yet pharmacological treatments remain elusive. Relapse can be prompted my multiple triggers, including drug-associated cues and context, stress, or the drug itself (Epstein *et al*. 2006). Animal models of cocaine relapse have been used to identify biological mechanisms that are altered by repeated use of cocaine and involved in relapse. The most prevalent model is the extinction-reinstatement paradigm in which animals undergo daily intravenous cocaine self-administration. During this time an operant response yields drug delivery and discrete drug-paired cues (e.g. stimulus light+tone). The self-administration period is followed by daily extinction training during which time the instrumental response made to obtain drug no longer delivers drug or cues. As extinction training typically takes place in the self-administration context, the drug-associated context is extinguished as well. Reinstatement of cocaine-seeking can then be induced by drug-associated cues, stress or the drug itself and is defined as the resumption of an extinguished response.

Using the extinction-reinstatement model, glutamate efflux increases in the NA core during cocaine-, cue-, and cocaine+cue-primed reinstatement of cocaine seeking (McFarland, Lapish & Kalivas 2003; Trantham-Davidson *et al*. 2012; Smith *et al*. 2017; Logan, LaCrosse & Knackstedt 2018)(Stennett, Padovan-Hernandez & Knackstedt 2019). This release is necessary for reinstatement as antagonism of post-synaptic glutamate receptors and stimulation of pre-synaptic autoreceptors prevents the reinstatement of cocaine-seeking (Scofield *et al*. 2016). Furthermore, compounds such as the beta-lactam antibiotic ceftriaxone which attenuate this NA core glutamate efflux also attenuate reinstatement of cocaine seeking (Trantham-Davidson *et al*. 2012). Ceftriaxone ameliorates several cocaine-induced NA core adaptations and consistently attenuates cocaine relapse across different relapse primes (Knackstedt, Melendez & Kalivas 2010a; Fischer, Houston & Rebec 2013; Bechard *et al*. 2018; Bechard & Knackstedt 2019).

The clinical treatment of cocaine use disorder rarely involves explicit instrumental extinction training. More often, treatment involves removing the person from their drug-associated context as they enter a drug-free period. Subsequent re-exposure to drug-associated contexts (with or without drug-paired cues) can prompt relapse. Thus, a model in which animals self-administer cocaine and then enter a drug-free abstinence period without instrumental extinction may have increased translational relevance. Relapse in this paradigm can be triggered by re-exposure to the drug-associated context alone or the context in combination with discrete drug-paired cues. When relapse is prompted by re-exposure to the context+cues after abstinence, this model is termed “incubation of cocaine craving model” because seeking increases with the length of abstinence. This is true after both extended (6 hr/day) and short (2-3 hr/day) access to cocaine self-administration, with increases in relapse seen after 15-45 days of abstinence (Grimm *et al*. 2001; Hollander & Carelli 2007; Lee *et al*. 2013; Cameron *et al*. 2019). In the present work, seeking was assessed on Day 15 of abstinence, but not on Day 1, and thus here the dependent variable (seeking on Day 15) is not referred to as “incubated cocaine seeking”. However, as in the “incubation” model, relapse is primed by cocaine-associated cues and context after a period of abstinence without extinction training.

Because extinction training produces distinct changes in the NA core which interact with medications to reduce relapse, it is important to utilize more than one model of relapse (Knackstedt *et al*. 2010b; Knackstedt, Trantham-Davidson & Schwendt 2014). Furthermore, the neural pathways underlying these types of relapse, although overlapping, are specific to their eliciting stimuli and altered by post-drug experience (McFarland & Kalivas 2001; Fuchs *et al*. 2005; Fuchs, Branham & See 2006; Knackstedt *et al*. 2010b). Brain regions mediating cocaine relapse include the infralimbic (IL) and prelimbic (PL) regions of the medial prefrontal cortex (mPFC), nucleus accumbens (NA) core and shell, and ventral tegmental area (VTA). Other nuclei that project to the NA, such as the basolateral amygdala (BLA), are activated during cue- and cocaine-primed relapse (McFarland & Kalivas 2001; Stefanik & Kalivas 2013), but not context-induced relapse in the absence of extinction training (Fuchs *et al*. 2006).

We propose that the identification of brain regions that mediate relapse after different post-drug experiences is essential for the development of pharmacotherapies to reduce relapse. To that end, the present set of studies tested the ability of ceftriaxone to attenuate context+cue-primed relapse after abstinence (incubation of cocaine seeking). We also quantified glutamate efflux in the NA core during relapse, with the hypothesis that ceftriaxone would attenuate relapse and glutamate efflux. In order to uncover the neurocircuitry engaged during context+cue-primed relapse and its interaction with ceftriaxone, we assessed expression of the immediate early gene Fos in regions that send glutamate projections to the NA core. In a separate experiment, we similarly trained rats after infusing the retrograde tracer cholera toxin B (CTb) into the NA core. CTb labels neurons that project to the NA core and overlap with Fos expression labels neurons projecting to the NA core active during relapse. Thus, we provide a novel neural map of the regions involved in cocaine relapse following context+cue-primed relapse, finding that attenuated cocaine-seeking is accompanied by reduced activation of NA core-projecting PL neurons.

## MATERIALS AND METHODS

### Animals

Male Sprague-Dawley rats (250-275 g) were acquired from Charles River LLC (NC, USA). Experiment 1 used 20 rats and Experiment 2 used 17 rats. Rats were housed individually on a 12 h reverse light cycle in a temperature-controlled vivarium, with all behavioral testing conducted in the dark phase. Rats were food restricted throughout, receiving 20 g food per day. All procedures were approved by the University of Florida’s Institutional Animal Care and Use Committee and followed the Guidelines of the Care and Use of Laboratory Animals.

### Drugs

Cocaine HCl was donated by the NIDA controlled substances program (RTI, Research Triangle, NC) and dissolved in 0.9% physiological saline as a 4 mg/mL solution. Ceftriaxone (Nova Plus) was dissolved in 0.9% physiological saline vehicle and administered intraperitoneally (IP) at 200 mg/kg in 1 mL/kg.

### Surgeries

Rats underwent surgical implantation of jugular catheters and microdialysis cannulae. Rats were anesthetized with IP ketamine (87.5 mg/kg) and xylazine (5 mg/kg). One end of the catheter (SILASTIC tubing, ID 0.51 mm, OD 0.94 mm, Dow Corning, Midland, MI) was implanted into the jugular vein. The other end exited from an incision on the rats’ back and was attached to a harness (Instech, Plymouth Meeting, PA) containing a cannula (Plastics One, Roanoake, VA) that could be connected to tubing for intravenous (IV) drug delivery. For Experiment 1, immediately after catheter implantation, unilateral microdialysis guide cannula were implanted above the NA core (AP +1.2 mm, ML +2.5 mm, DV −5.0 mm) as in our previous work on this topic (Trantham-Davidson *et al*. 2012; Stennett *et al*. 2019). In Experiment 2, rats instead received intra-NA core Alexa Fluor™ 488 conjugated CTb (Thermo Fisher Cat. No. C-34775). CTb was administered in five injections (50 nL/injection) with a Nanoject II (Drummond Scientific, Broomall, PA, USA) into the NA core using the same A/P and M/L coordinates as the microdialysis cannulae, such that injections were delivered to span the region sampled by the microdialysis probe in Experiment 1 (5 injections at 5 locations between −7.0 and −5.5 DV). Postoperative care consisted of 2 days of antibiotic (Cefazolin, 100 mg/mL, IV) and the analgesic Carprofen (2.5 mg/kg), and daily flushing of the catheters using heparinized saline (0.2 mL of 100 IU, IV) that continued throughout the self-administration period. Catheter patency was tested at regular intervals with sodium brevitol (10 mg/mL, IV; Eli Lilly, Indianapolis, IN, USA).

### Cocaine self-administration and relapse test (Experiment 1 and 2)

Rats were trained to press a lever to receive an infusion of cocaine (1.0 mg/kg per 0.10 μL infusion) in a two-lever operant chamber (30 × 24 × 30 cm; Med Associates, St. Albans, VT, USA). Drug delivery was accompanied by presentation of discrete cues: a stimulus light over the active lever and a 2900 Hz tone for the duration of the infusion (5 sec). Presses on the active lever yielded drug and cues on a fixed ratio-1 schedule of reinforcement, whereas presses on the inactive lever were recorded but were not reinforced. Infusions were accompanied by a time-out period (20 sec), during which time lever presses were recorded but not reinforced. Rats self-administered cocaine for 2 hr/day for a minimum of 12 days, with the additional criteria of 9 or more infusions/day in the last 6 days of self-administration and no more than 25% variability in infusions/active lever presses in the last 3 days. Subsequently, animals entered a period of abstinence where they were handled and weighed daily for 2 weeks but not put back into the operant chamber. During the last six days of this abstinence period, rats were given a daily injection of ceftriaxone (Cef; 200 mg/kg IP) or its vehicle (Veh; 0.9% physiological saline). Rats were then re-exposed to the context for a relapse test (100 min) during which time presses on the active lever yielded the drug-associated cues but no drug. The length of the test was chosen based on our recent assessment of the same dependent variables (Fos and glutamate efflux) with a similar timed test (Stennett *et al*. 2019). See Fig. 1a for timeline.

**Figure 1.**
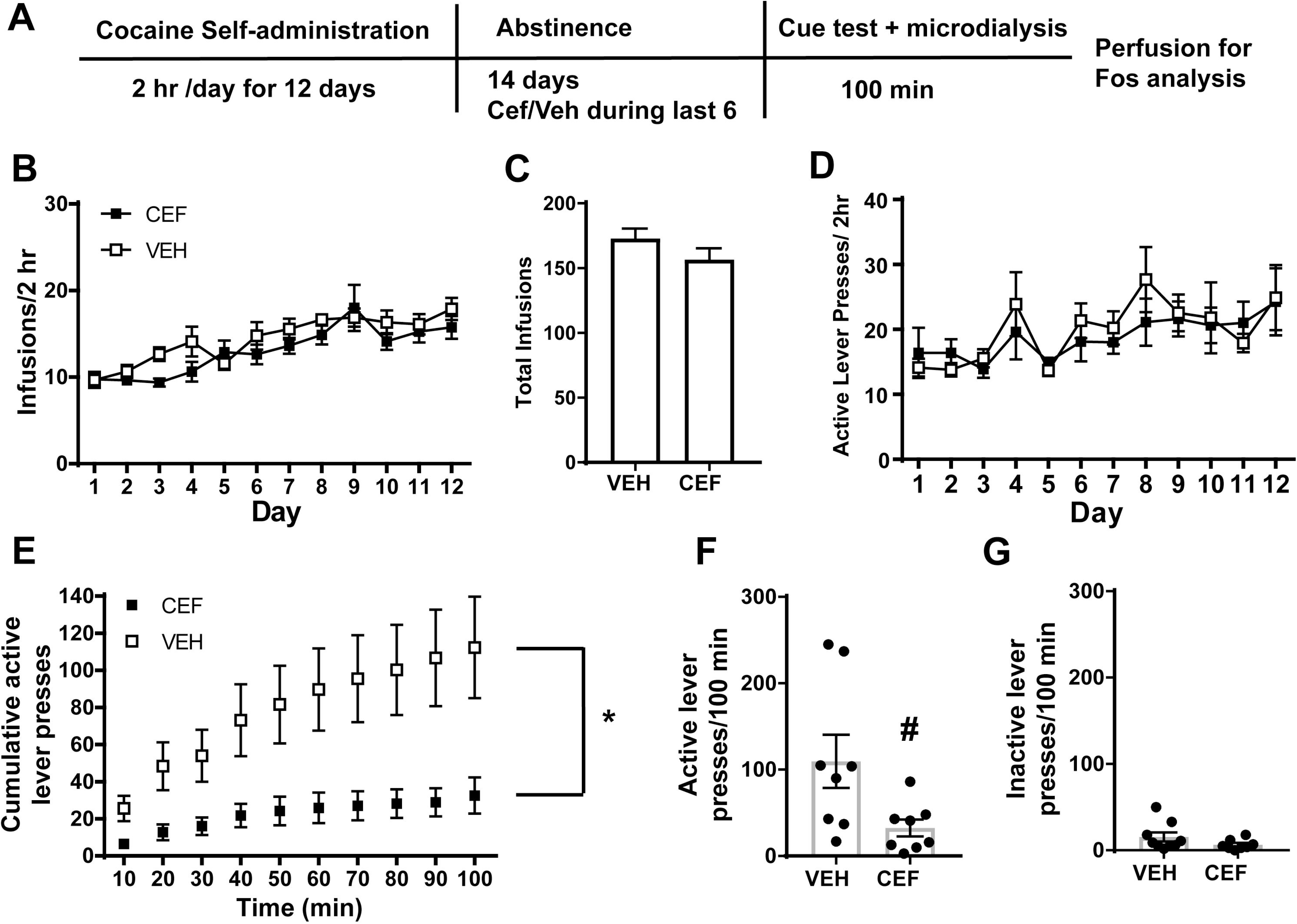
Ceftriaxone attenuates cue+context-primed relapse of cocaine seeking after abstinence. **A.** Timeline of Experiment 1. Infusions attained during self-administration differed by group (**B**), however total number of infusions attained was similar (**C**). **D.** There were no differences in active lever pressing during self-administration between rats that later received Cef/Veh during abstinence. **E.** Ceftriaxone reduced cumulative active lever pressing during relapse and **(F)** the total number of active lever presses during the test, with no effect on inactive lever pressing **(G).** * = p<0.05 main effect of Group. # = p < 0.05 compared to Veh.

### Experiment 1

Twenty rats self-administered cocaine as described above. On the night before the relapse test, rats were implanted with microdialysis probes and perfused with artificial spinal fluid (aCSF). Rats spent the night in their home cages adjacent to the operant chamber. The next morning, 2 hours of baseline samples were collected while rats were still in the home cage. Rats were then placed inside the operant chambers for the cue-primed relapse test and samples collected every 10 minutes. Lever presses were recorded in 10-minute bins. After 100 minutes, rats were given a lethal dose of pentobarbital (100 mg/kg IP) and transcardially perfused. Brains were harvested and stored at −80°C until further processing.

Glutamate concentration in microdialysis samples was quantified using isocratic high-performance liquid chromatography (HPLC) with electrochemical detection (Thermo Fisher Scientific). O-pthalaldehyde (Sigma-Aldrich, USA) derivatized samples prior to injection onto a CAPCELL PAK C18 column (5 μm, 2.0 mm I.D. X 50 mm; Shiseido). Mobile phase consisted of 100 mM Na_2_HPO_4_, 10% v/v methanol, and 3.5% v/v acetonitrile (pH 6.5). The amount of glutamate in each sample was quantified by comparing computer-integrated peak areas of samples with those of known glutamate standards.

Brains were sliced and stained to assess the effects of Cef on Fos expression in regions known to regulate relapse to cocaine-seeking after abstinence or extinction, namely the IL and PL regions of the mPFC, NA core and shell, BLA, VTA, and paraventricular nucleus of the thalamus (PVN). Fos was not quantified in the entire NA core, but only in the region sampled by microdialysis probes in this study and in our previous studies. See Fig. 3A for regions of interest. Free-floating sections (30 μM) were quenched in H_2_O_2_ (0.3% in PBS), blocked in normal donkey serum in PBST (2%) and transferred to rabbit anti-Fos primary antibody (1:10000; EMD Millipore) in PBST and normal donkey serum for 16 hr at RT. Sections were washed and transferred to donkey anti-rabbit secondary antibody (1:500; Jackson Immuno) for 2 hr before detection with ABC (Vectastain ABC Kit, Vector Laboratories) and visualization with DAB substrate (Vector Laboratories). Images were acquired with a Tucsen monochrome CCD camera attached to an Olympus BX51 fluorescent microscope and quantified using Image J software (NIH). Between 2 and 4 sections/region were acquired and analyzed and the average number of Fos+ cells/mm^2^ within each region were calculated for each rat. Slices would occasionally be missing from one brain region or mis-stained, resulting in different n/group for each brain region.

### Experiment 2

Following surgery to implant catheters and administer intra-Nac CTb, 17 rats underwent cocaine self-administration and relapse testing as described above. Rats were perfused immediately after the test. Sections of tissue (15 μm) were mounted onto a Superfrost plus slides (Fisher Scientific) and the perimeter was sealed with a hydrophobic barrier pen (Vector Laboratories). Sections were then washed in 0.3% PBST and blocked in glycine buffer (1 hr) followed by blocking in normal goat serum (10%) in PBST (2 hr). Rabbit anti-fos primary (1:1000) was applied and left to sit overnight (16 hr) at RT. The following day, slices were washed and incubated in Alexa Fluor-594 secondary (goat anti-rabbit) for 2 hr and coverslipped with ProLong Gold antifade reagent mounting medium.

We captured fluorescent images at 40x magnification using a Tuscen CMOS camera mounted on an Olympus BX51 microscope. Excitation filters captured DAPI, CTb, and Fos images at 358 nm, 488 nm, and 594 nm, respectively. Single-channel images were overlayed. At least 4 composite images (2 from each hemisphere) were obtained from each rat from each region (PL and IL) and counted (4.20 – 2.52 mm anterior to bregma, Paxinos & Watson, 2007). Composite images were imported into NIH Image J software (Schneider et al., 2012). Reviewers blind to experimental conditions quantified CTb^+^ and Fos^+^ cells using a Cell Counter plugin offered by NIH (available at https://imagej.nih.gov/ij/plugins/cell-counter.html). DAPI was used to confirm the presence of cell nuclei. Fos expression was established when corresponding fluorescent protein was enveloped within or abutting the cell nucleus, presumably within the cytoplasm (Bussolino *et al*. 2001; Caputto, Cardozo Gizzi & Gil 2014). The ratio of Fos^+^ cells expressing CTb was calculated for each rat, as was the converse (number of CTb^+^ cells expressing Fos).

### Statistical Analysis

Analyses were conducted using GraphPad Prism (8.0). Self-administration data were analyzed using a Repeated Measures (RM) ANOVA, with Time as the within-subjects factor and group (Cef/Veh) as the between-subjects factor. The number of infusions, active lever presses, and inactive lever presses were compared between groups later receiving Cef/Veh during abstinence to ensure there were no pre-treatment difference in cocaine intake prior to treatment. The effect of ceftriaxone on active lever pressing during the relapse test was examined in two ways for Experiment 1. First, responses were summed into 10 minute bins so that the cumulative number of active lever presses across the relapse test could be: 1) compared between groups using a RM ANOVA with Time and Group as factors and 2) correlated with glutamate efflux during these 10-min bins. Second, the total number of active and inactive lever presses during the relapse test was assessed for differences between groups using an independent samples t-test. The relationship between the total number of active lever presses during relapse and total Fos expression within a brain region was examined using Pearson’s correlation. For Experiment 2, only total number of active and inactive lever presses were compared between groups. The amount of co-labeled Fos and CTb was compared between groups with independent samples t-tests. Glutamate concentrations were compared between groups in two ways, both the raw values and the percent change in were compared using RM ANOVA with Time and Group as factors.

## RESULTS

### Experiment 1

This experiment ran 20 rats through cocaine self-administration. Two rats were excluded for not meeting self-administration criteria. One rat from the vehicle-treated group exceeded 2 standard deviations from the mean for active presses during the relapse test and was excluded from analysis. No Group × Time interaction was found for infusions across the 12 days of self-administration (Fig. 1B). However, a main effect of Group was detected [F_(1,180)_=7.784, p=0.0058]. To confirm that the total number of infusions attained during self-administration did not differ between rats later assigned to receive Cef or Veh, a t-test was conducted, finding no significant difference (Fig. 1C). There was no main effect of Group nor a Group × Time interaction for active lever presses during self-administration (Fig. 1D). The number of infusions and active lever presses escalated over time in both groups [infusions: F_(11,180)_=9.461; p<0.0001; active lever presses: F_(11,180)_=2.586; p=0.0045]. Inactive lever presses did not differ by Time or Group. Cef reduced cumulative active lever presses during the cue+context test, as there was a main effect of Group [F_(1,150)_=60.18, p<0.0001; Fig. 1E]. There was no Group × Time interaction, likely because cumulative active lever presses increased similarly over time for both groups [F_(9,150)_=3.3, p =0.001]. The total number of active lever presses (Fig. 1F) during relapse was reduced by Cef [t_(14)_=2.388, p=0.0316]. No difference in cumulative (not shown) or total (Fig. 1G) inactive lever presses was found.

Five rats (2 Cef and 3 Veh) experienced microdialysis probe failure and did not contribute to the microdialysis data but still underwent relapse testing for Fos analysis. This yielded 6 Cef- and 6 Veh-treated rats for which microdialysis data were collected. During the relapse test, glutamate efflux (% baseline) was reduced by Cef. A Group × Time interaction was detected [F_(13,130)_=1.99, p=0.0263], as well as a significant effect of Group [F_(1, 10)_=10.96, p=0.0079], but not Time (Fig. 2A). When examining raw glutamate values (Fig. 2B), the same results were obtained [Interaction: F_(13,130)_=3.094, p=0.0005; main effect of Group: F_(1,10)_=7.158, p= 0.0233]. Post-hoc tests conducted on both raw and % baseline glutamate found that efflux increased from baseline only in vehicle-treated rats, and group differences were detected at several times. The mean percent change in glutamate efflux was calculated for each group for each 10-minute bin and was found to correlate with mean cumulative active lever pressing (r=0.7355, n=20, p=0.0002; Fig. 2C). Placement of dialysis probes is noted in Fig. 2D.

**Figure 2.**
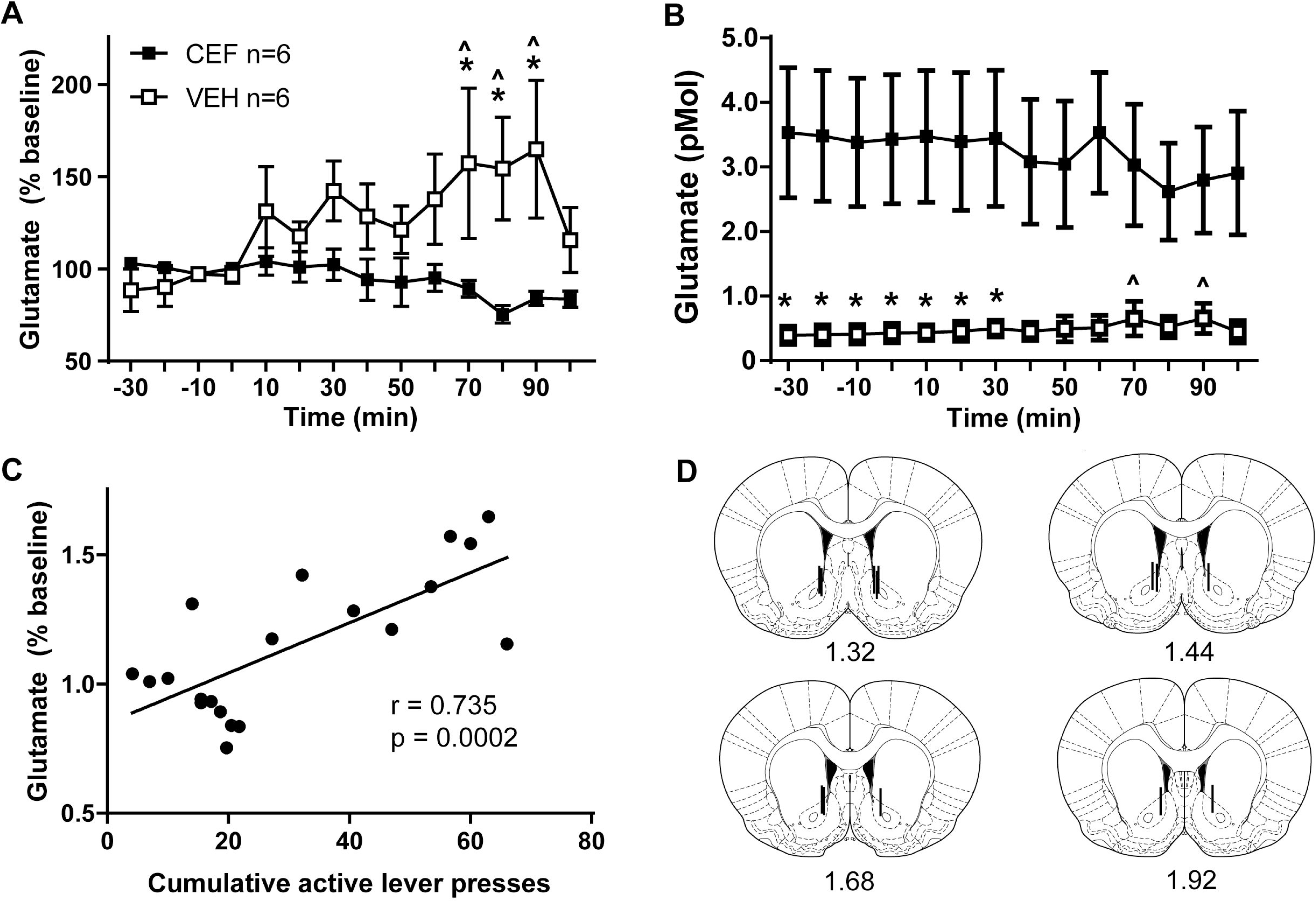
Ceftriaxone increases baseline glutamate levels while decreasing glutamate efflux during relapse. **A.** During relapse, vehicle-treated rats display increased glutamate (percent baseline levels) relative to both their own baseline and to the ceftriaxone-treated group. **B.** Examining raw glutamate levels shows the same pattern, as well as an effect of ceftriaxone on increasing glutamate levels prior to the relapse test. **C.** The changes in glutamate efflux during relapse are correlated with cumulative active lever presses during relapse testing. **D.** Placement of microdialysis cannulae. * = p<0.05 compared to Cef; ^ = p<0.05 compared to own baseline.

Relapse-induced Fos expression was reduced in Cef-treated rats in several brain regions: NA core (t_(11)_ =2.361, p=0.0377, Fig. 3B); PL (t_(12)_=2.335,p=0.0378, Fig. 3D); BLA (t_(14)_=3.032, p=0.0090, Fig. 3E); VTA (t_(12)_=3.365, p=0.0056, Fig. 3F). There was a trend for Cef to reduce Fos expression in the NA shell (t_(12)_=2.128, p =0.054, Fig. 3C). No group differences were observed in the IL or PVN (not shown). Total active lever presses during relapse positively correlated with Fos expression in the PL (r=0.5824, p=0.0289), BLA (r=0.5029, p=0.0471), and VTA (r=0.7139, p=0.0041). No such relationship was found for the NA core, NA shell, IL or PVN. A control group not receiving cocaine was not used for this study, as we have reported minimal (<5) Fos+ cells per region in this condition (Stennett *et al*. 2019).

**Figure 3.**
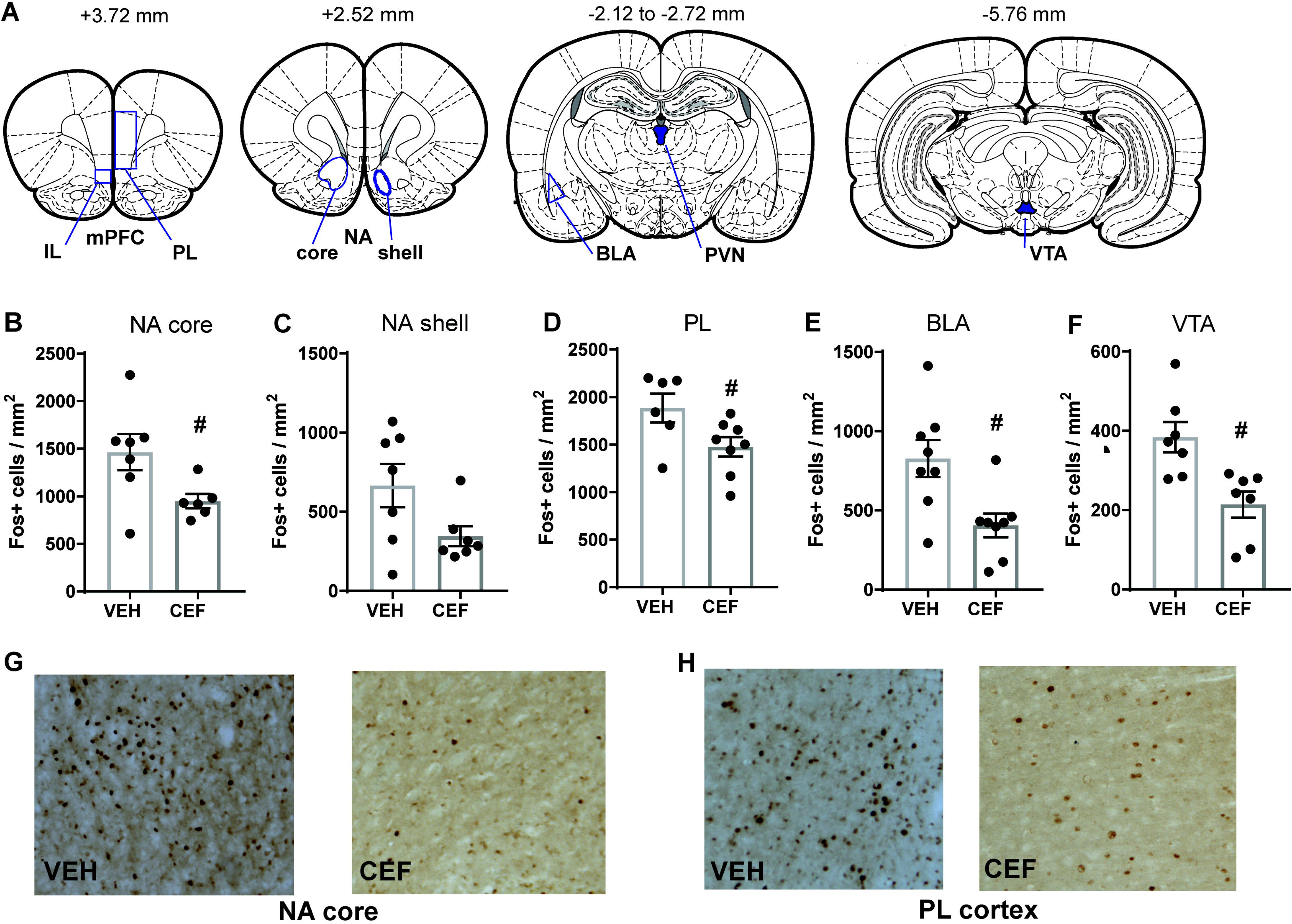
Post-relapse Fos expression is reduced by ceftriaxone in several brain regions. **A.** Sites of Fos analysis. Fos expression after relapse testing in the NA core (**B**), NA shell (**C**), PL (**D**), BLA (**E**) and VTA **(F)**. Fos expression was reduced by ceftriaxone in the NA core, PL, BLA and VTA. **G.** Sample images from the NA core and **(H)** PL cortex from Cef and Veh-treated rats. # = p<0.05 compared to Veh.

### Experiment 2

In Experiment 2, 1 rat experienced catheter failure, and 3 additional rats were eliminated for failing to meet self-administration criteria. One rat was eliminated from data analysis post-hoc due to CTb injection outside of the NA core, leaving 5 vehicle-treated rats and 7 ceftriaxone-treated rats included in the analysis. Self-administration behavior did not differ between Cef and Veh groups as shown by the number of infusions and active lever presses, although both increased over Time [infusions: F_(11,121)_=8.897; p<0.0001; active lever presses: F_(11,121)_=2.234; p=0.0167; see Fig. 4A,B]. Inactive lever presses were low throughout self-administration, with no group differences (not shown). For this experiment, we assessed the total number of lever presses during the 100 minute relapse test, finding a significant reduction in active lever presses by Cef [t_(10)_=2.81, p=0.0185; Fig. 4C]. Inactive lever presses during the test were unaffected by Cef (Fig. 4D).

**Figure 4.**
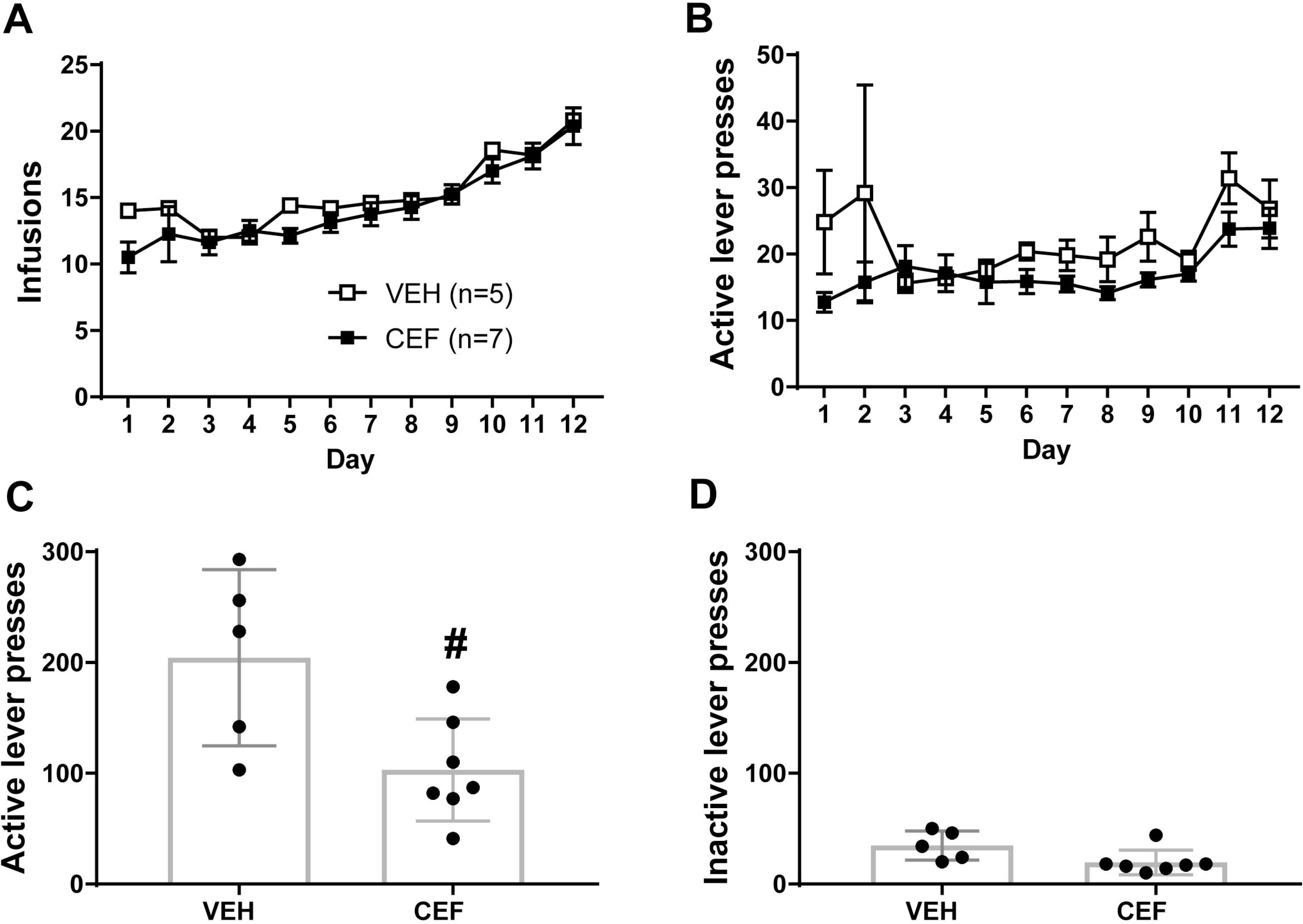
Ceftriaxone attenuates cue+context-primed relapse of cocaine seeking after abstinence. **A.** Infusions and (**B**) active lever presses did not differ between groups later assigned to receive Cef/Veh. **C.** Cef attenuated active lever pressing during the relapse test, with no effect on inactive lever presses (**D**). # = p<0.05 compared to Veh.

Following injection in the dorsal medial region of the NA core where microdialysis probe placement occurred in Experiment 1 (Fig. 2D and 5A), and our previous publications (Trantham-Davidson *et al*. 2012; Stennett *et al*. 2019), CTb was observed in the IL and PL, BLA, PVN, and VTA (Fig. 5B–C). In the PL, 74% of CTb-labeled cells were also Fos+ in vehicle-treated rats, indicating that NA core-projecting neurons in this region are highly active during a cue+context primed relapse test. This number was reduced in ceftriaxone-treated rats [t_(9)_=2.41, p=0.0393, Fig. 5D]. Similar numbers of Ctb-labeled cells were activated in the IL of vehicle-treated rats, but ceftriaxone did not alter this measure [t(9)=1.93, p=0.08, Fig.5F]. Conversely, only 36.4% of Fos+ cells were also labeled with CTb in the PL of vehicle-treated rats, indicating that the majority of relapse-active cells in this brain region do not project to the NA core. Ceftriaxone reduced this number (t_(9)_=2.704, p=0.0242, Fig. 5E]. The number of Fos+ cells also labeled with CTb was less than 30% in the IL and did not differ between Cef and Veh groups (Fig. 5G).

**Figure 5.**
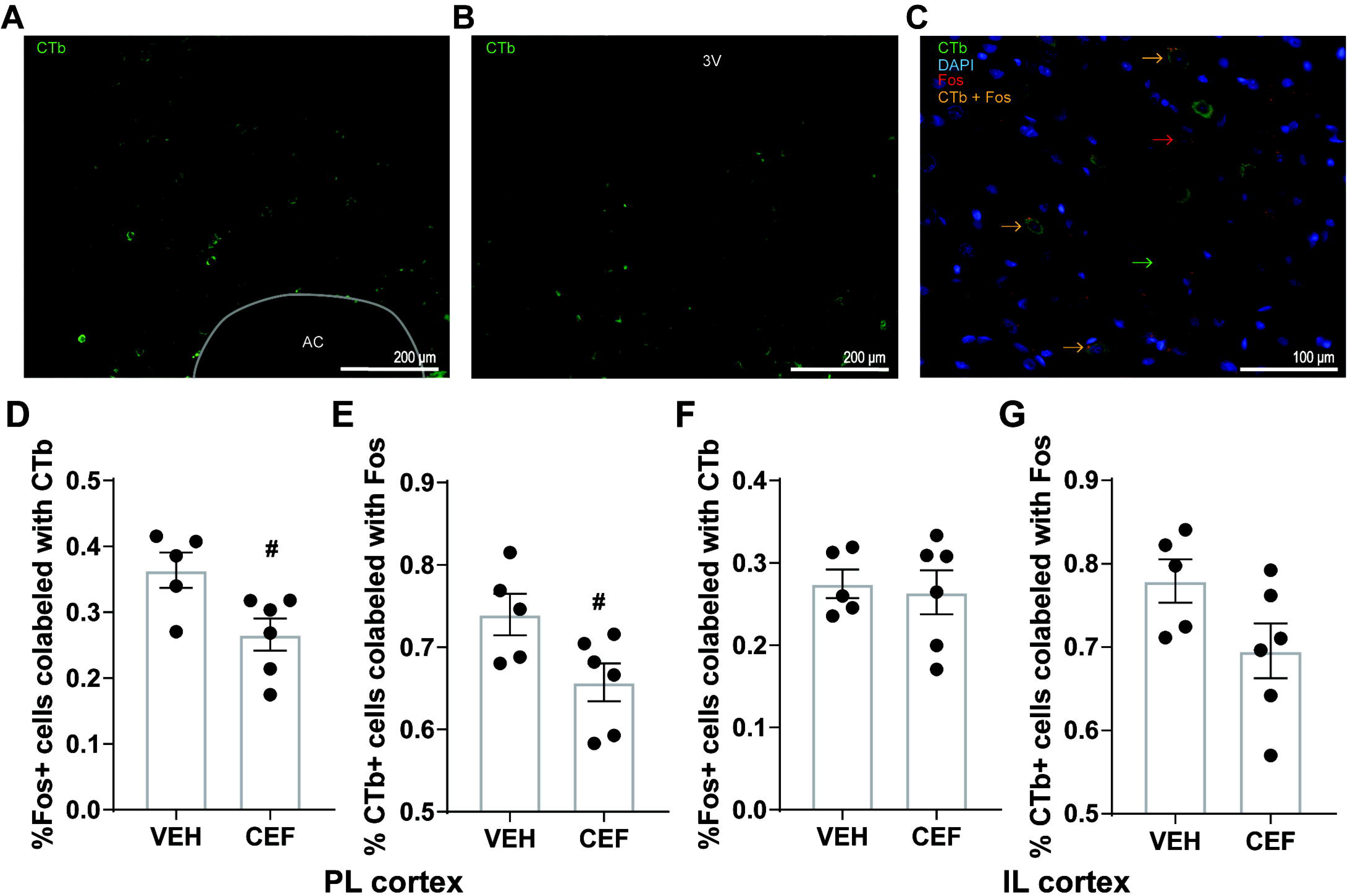
**A.** Spread of CTb within nucleus accumbens core, surrounding the anterior commissure (AC), taken at 20x magnification. **B.** Spread of CTb within the paraventricular nucleus, beneath the third ventricle (3V); 20x magnification. **C.** Triple-labeled image (40x magnification) depicting co-localization of Fos and CTb within the infralimbic prefrontal cortex (PFC); DAPI was used to stain nuclei and confirm presence of cell bodies. **D.** Percentage of Fos+ cells that also express CTb within prelimbic PFC. **E.** Percentage of CTb+ cells that also express Fos within prelimbic PFC. **F.** Percentage of Fos+ cells that also express CTb within infralimbic PFC. **G.** Percentage of CTb+ cells that also express Fos within infralimbic PFC. # = p<0.05 compared to Veh.

### Discussion

Here we show for the first time that glutamate efflux in the NA core accompanies drug-seeking in a context+cue-primed test of relapse after abstinence without extinction training. While we did not assess a time dependent increase in seeking in our model from early to late abstinence, incubated cocaine seeking is observed after 14 days of abstinence from 2 hr/day cocaine self-administration (Cameron *et al*. 2019). Glutamate release is attenuated by subchronic treatment with ceftriaxone and accompanied by reduced Fos activation in the NA core itself, and regions that project to the NA core, such as BLA, VTA, and the PL cortex. The attenuation of Fos expression by ceftriaxone implies that such expression is related to relapse. These findings indicate that like context-primed relapse after abstinence, and cue, cocaine and cue+cocaine primed reinstatement after extinction, incubation of cocaine seeking is accompanied by glutamate efflux in the NA core that can be attenuated by ceftriaxone (McFarland *et al*. 2003; LaCrosse, Hill & Knackstedt 2016; Smith *et al*. 2017; Stennett *et al*. 2019). To determine if activity is reduced in neurons sending glutamate projections to the NA core, we then labeled such neurons with the retrograde tracer CTb, finding that ceftriaxone-treated rats showed reduced activation in PL and not IL neurons that project to the NA core.

We previously reported that subchronic ceftriaxone attenuates context-primed relapse and glutamate efflux after abstinence, but glutamate efflux still increases from baseline (LaCrosse *et al*. 2016). It is possible that extinction training interacts with ceftriaxone to normalize glutamate efflux, potentially due to ceftriaxone’s ability to increase GLT-1 in the PFC after extinction training (Sari et al. 2009), but not abstinence (LaCrosse et al. 2016). However, the current data show that glutamate efflux is suppressed by ceftriaxone treatment in the absence of extinction training when relapse is primed by both context and drug-associated cues. We propose that this difference arises from the BLA and PL involvement in relapse when discrete drug-associated cues prime relapse. Inactivation of these regions has no effect on relapse primed by context alone after abstinence (Fuchs *et al*. 2006), but attenuates reinstatement in a cue-primed test after extinction (Stefanik *et al*. 2013; Stefanik & Kalivas 2013). Here ceftriaxone reduced overall Fos expression in the BLA and PL, and Fos expression in NA core-projecting PL neurons. Furthermore, Fos expression in the BLA correlates both with active lever pressing during relapse and the amount of glutamate release (AUC) during relapse. Thus, in this model, our data implies that ceftriaxone attenuates NA core glutamate release during context+cue-primed relapse via suppressing activity of BLA and PL glutamate neurons.

In addition, our data implicate differential effects of ceftriaxone in the subregions of the mPFC. The mPFC’s discrete subregions mediate different aspects of drug-seeking, with the PL projections to the NA core promoting cue-primed drug-seeking (Stefanik *et al*. 2013), and IL projections to the NA shell promoting extinction learning and inhibition of cocaine-seeking (Ongür & Price 2000; West *et al*. 2014; Augur *et al*. 2016). Moreover, functionally distinct neural populations in the PL and BLA are evidenced by Fos expression in neurons projecting to the NA core, but not shell, upon cue-primed reinstatement of cocaine seeking (McGlinchey *et al*. 2016). We are the first to examine the relationship between attenuation of cocaine relapse with the activity of NA core-projecting PL neurons. Notably, approximately 75% of NA core-projecting PL neurons expressed Fos in the vehicle group. These number were reduced by ceftriaxone treatment, as was the percent of Fos+ cells also expressing CTb. These results implicate inhibition of neurons projecting from PL to NA core in the ability of ceftriaxone to attenuate context+cue-primed relapse after abstinence, consistent with a role for this pathway in mediating cue-primed reinstatement after extinction (Stefanik *et al*. 2013). Pharmacological inhibition of the PL not only attenuates cocaine-primed reinstatement after extinction, but also glutamate efflux in the NA core during relapse (McFarland *et al*. 2003). Blocking dopamine receptors in the PL and glutamate receptors in the contralateral NA core attenuates cue-induced reinstatement of cocaine-seeking after extinction, and Fos expression in NA core-projecting PL neurons correlates with cue-primed reinstatement (McGlinchey *et al*. 2016). Thus, the PL alone and PL-NA core pathway is critical for relapse to cocaine seeking after both extinction and abstinence.

After extended access cocaine self-administration and 30 days of withdrawal, incubated cocaine seeking is suppressed by pharmacological inactivation of the IL cortex (Koya *et al*. 2009). However, optogenetic stimulation of IL-NA shell projections suppresses relapse during a context+cue test of incubated cocaine-seeking (Cameron *et al*. 2019). When incubated cocaine seeking is observed after 14 days of withdrawal from 2 hr/day cocaine self-administration, activity in NA shell-projecting IL neurons decreases prior to execution of the drug-seeking response (Cameron *et al*. 2019). The magnitude of activity of IL-NA shell projections immediately after execution of the drug-seeking response during the relapse test is inversely related to the number of subsequent drug-seeking responses made. Thus, the lack of an effect of ceftriaxone on total IL Fos expression and Fos expression in IL-NA core projecting neurons is consistent with the idea that these neurons are not involved in driving the drug-seeking response during a context+cue-primed test after abstinence. We previously reported that IL Fos expression was induced during a cocaine+cue-primed reinstatement test following extinction training, and that ceftriaxone attenuated such expression (Stennett *et al*. 2019). The difference between these results may arise from different primes or post-cocaine training regimen (abstinence vs. extinction).

Relapse-induced Fos expression was reduced in other brain regions that project to the NA core, namely the BLA and VTA; such activity correlated with active lever pressing during relapse. Future work will examine whether this reduction in activity occurred in neurons projecting to the NA core. In addition to the well-characterized dopamine projections from VTA to NA core that regulate motivated behavior, recently identified subpopulations of VTA neurons that co-release GABA and glutamate have been shown to have distinct roles in reinforcement, motivation and learning (Brown *et al*. 2012; Morales & Margolis 2017; Guzman *et al*. 2018). Fos expression arises from signaling through both glutamate and dopamine receptors (Young, Porrino & Iadarola 1991; Wang 1998), leaving open the possibility that the reduction in VTA Fos expression after ceftriaxone underlies reduced Fos expression in regions receiving dopaminergic projections from the VTA. Future studies should identify which VTA neuronal subpopulations exhibit Fos expression upon relapse and the projection targets of such neurons.

Interestingly, while ceftriaxone-treated rats displayed reduced Fos in the NA core overall, such activity did not correlate with active lever pressing during relapse. The overall reduction in NA core Fos may be due to reduced glutamate (and possibly DA) release arising from PL, VTA, and BLA. The lack of correlation may arise from Fos expression in the NA core during relapse occurring in both D1- and D2-positive neurons, which have opposing roles in regulating cocaine-seeking prompted by cues (Bock *et al*. 2013).

In conclusion, the present work finds for the first time, glutamate efflux accompanies incubated cocaine seeking after abstinence and implicates the PL-NA core projection in this type of relapse. The results of the Fos mapping studies are consistent with the hypothesis that ceftriaxone not only alters glutamate homeostasis within the NA core (Knackstedt *et al*. 2010a; LaCrosse *et al*. 2017; Bechard *et al*. 2018), but also affects several regions that send glutamatergic projections to the NA core to reduce both relapse and the glutamate efflux that accompanies it.

## Acknowledgements

This work was funded by NIH grants DA033436 and DA045140 awarded to LAK. LAK and ARB designed the studies. ARB and CNL conducted microdialysis and HPLC procedures. ARB, HB, JM, VLH and YPH contributed to imaging and analysis of Fos and Fos+CTB. ARB, CNL, JM and LAK analyzed the data and wrote the manuscript.

